# Phylogenetic, genomic, and morphological characterization of *Rickettsia senegalensis* sp. nov., a novel *Rickettsia* species detected worldwide

**DOI:** 10.64898/2026.05.02.721834

**Authors:** Clément Labarrere, Cheikh Tidiane Houmenou, Pierre-Edouard Fournier, Florence Fenollar, Oleg Mediannikov

## Abstract

Members of the genus *Rickettsia* are widespread obligate intracellular bacteria with high medical and veterinary relevance. While *Rickettsia senegalensis* was previously isolated from cat fleas, *Ctenocephalides felis*, in Senegal, a comprehensive polyphasic characterization is required for its formal taxonomic validation. This study aims to provide the first detailed phenotypic, ultrastructural, and genomic characterization of the type strain PU01-02ᵀ. Cultivation assays demonstrated that strain PU01-02ᵀ grows aerobically in XTC-2, SF9, and LD652 cell lines at 28 °C in a CO₂-free atmosphere. Ultrastructural examination via scanning electron microscopy using a backscattered electron detector confirmed its free cytoplasmic localization and a typical rickettsial morphology characterized by rod-shaped cells approximately 1.5 µm in length and 0.3 µm in width. Whole-genome sequencing yielded a 1.62-Mb genome with a G+C content of 33.2%, and three plasmids were detected. Genomic analyses confirmed its status as a distinct species, placing it within the transitional *Rickettsia* group, specifically inside the *Rickettsia felis* cluster. A review of epidemiological data revealed that rickettsial genes identical to those of *R. senegalensis* had already been identified in several hematophagous arthropods, including fleas and ticks parasitizing various hosts such as cats, dogs, opossums, and rodents worldwide. It has also been detected in cat tissues, suggesting a potential host-pathogen association. Based on these comprehensive characterization data, we formally propose *Rickettsia senegalensis* sp. nov. as a new species, with PU01-02ᵀ (= CSUR R184ᵀ = DSM 28250ᵀ) as the type strain.

**Repositories:** The genome sequence of *Rickettsia senegalensis* sp. nov. strain PU01-02^T^ has been deposited in GenBank under accession number JBVYTQ000000000, and the *rrs*, *gltA*, *ompB* and *sca4* gene sequences under accession numbers KF666476, KF666472, KF666470, KF666474, respectively. The plasmid accession numbers are PZ272915, PZ272916, and PZ272917, for pRS01, pRS02 and pRS03, respectively.

## INTRODUCTION

*Rickettsia* species (Rickettsiales, Rickettsiaceae) are Gram-negative, obligate intracellular bacteria responsible for various human diseases, including epidemic typhus [1]. To date, the genus *Rickettsia* comprises more than 30 species, 22 of which are pathogenic to humans [2, 3]. These species are classified into four groups based on their genomic, epidemiological, and clinical characteristics: the ancestral group (AG), the spotted fever group (SFG), the typhus group (TG), and the transitional group (TRG) [4, 5].

Several species within the TRG, such as *Rickettsia akari* and *Rickettsia australis*, are well-established human pathogens [6, 7]. More recently, increasing attention has been given to *Rickettsia felis* and closely related species within this group. *Rickettsia felis* has been detected in more than 20 different species of hematophagous arthropods worldwide, including fleas, *Anopheles* mosquitoes, lice, soft and hard ticks [8–10]. Non-hematophagous booklice *Liposcelis bostrychophila* were also found to harbour *R. felis* [11]. Considered an emerging *Rickettsia*, *R. felis* has been implicated in cases of fever reported on every continent except Antarctica. Studies have demonstrated that *R. felis* is a significant cause of febrile illnesses of unknown origin, accounting for up to 15% of cases in sub-Saharan Africa and reaching 19.6% in parts of Asia [12–16]. It seems that *R. felis*-associated fever is present in tropical and sub-tropical regions all over the world and sometimes may be quite prevalent. *Rickettsia felis* is the most extensively studied species within a cluster that comprises multiple arthropod-associated bacteria, most of which have not yet been isolated and characterized [17, 18]. However, the genetically close *Rickettsia asembonensis* species, recently described from fleas, was also implicated in human pathology [19, 20]. Consequently, characterizing other members of this *R. felis* cluster is critical, as they share similar arthropod vectors and geographic niches, yet their specific medical and veterinary significance remains largely unresolved.

In this context, this study describes *Rickettsia senegalensis,* another prominent member of the *R. felis* cluster. Isolated in 2012 from cat fleas, *Ctenocephalides felis*, collected in Dakar, Senegal, this species has since been detected multiple times in different regions worldwide (Figure 1) [21]. It has been identified in various hematophagous arthropods, including fleas parasitizing cats, dogs, opossums, and rodents (*C. felis, Ctenophalides orientis, Coptopsylla lamellifer, Xenopsylla skrjabini, Xenopsylla gerbilli, Echidnophaga gallinacea*), as well as in tick such as *Dermacentor nitens*, *Rhipicephalus microplus* (Table 1). Additionally, this *Rickettsia* has been detected in cat tissues in California, USA, suggesting pathogenicity for mammals. However, relying solely on molecular detection limits our understanding of its biology, physiology, and actual pathogenic potential. Obtaining and maintaining a viable, pure isolate remains a major bottleneck in rickettsiology, yet it is essential for performing comprehensive phenotypic, ultrastructural, and high-quality genomic characterizations.

**Figure 1.**
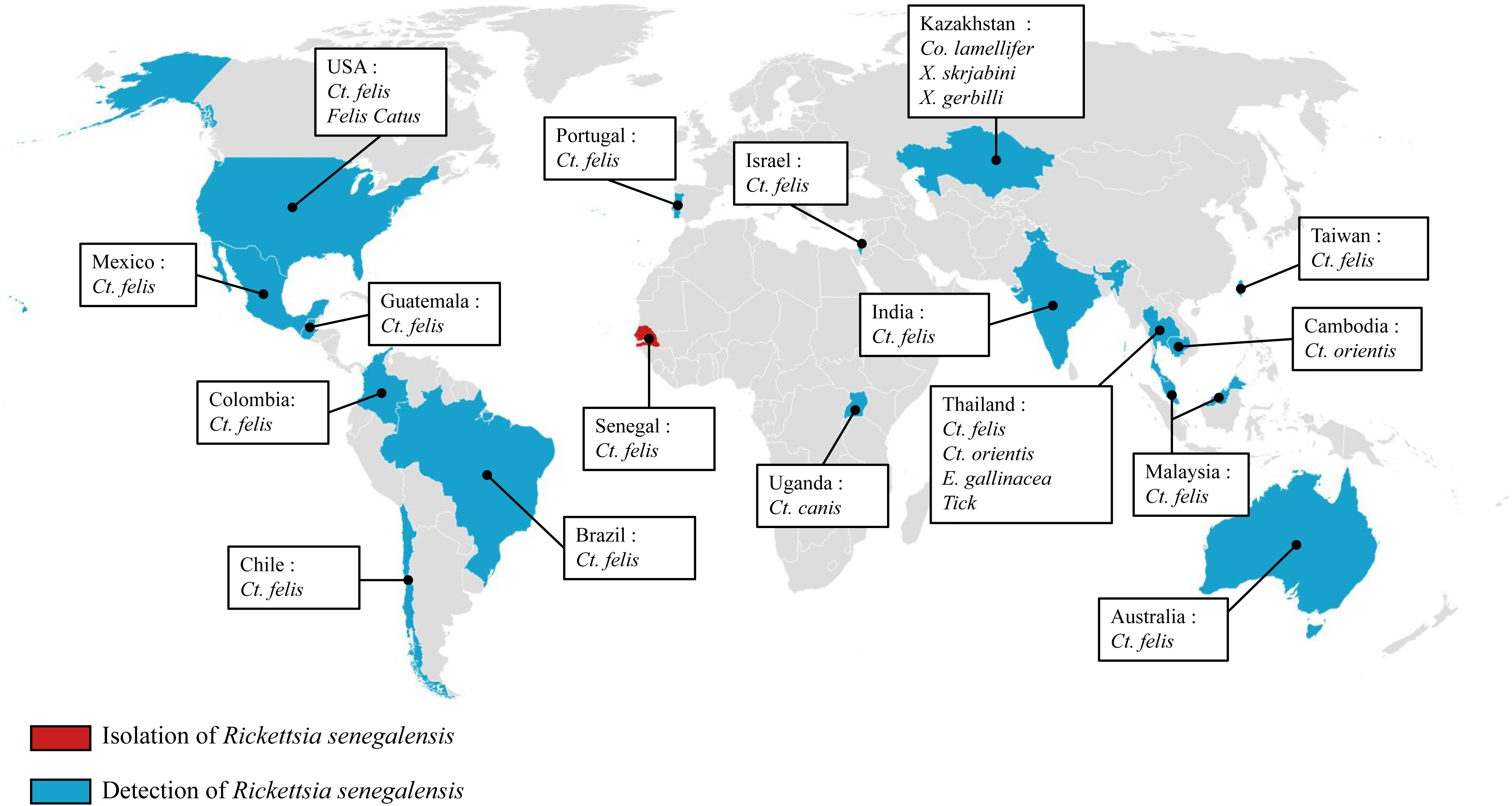
Global distribution of Rickettsia senegalensis and its hosts. Reported occurrences include fleas (Ctenocephalides felis, Ctenophalides canis, Ctenophalides orientis, Coptopsylla lamellifer, Xenopsylla skrjabini, X. gerbilli, Echidnophaga gallinacea), ticks, and domestic cats (Felis catus) across 14 countries, with isolates obtained from fleas (Ct. felis) in Senegal.

**Table 1.** Summary of the identification of *Rickettsia senegalensis* reported to date. The table includes information on the country, designation, date, collection source, host, the prevalence of samples from fleas, ticks and cats testing positive for *R. senegalensis* and corresponding references. ND: No data.

| Country | Designation | Date | Collection source | Host | Number of samples tested and prevalence | Refs |
| --- | --- | --- | --- | --- | --- | --- |
| Senegal | Strain PU01-02<sup>T</sup> | 2012 | Cat | *C. felis* | 5 positive fleas / 29 fleas | [21] |
| California, USA | Isolate Cf US 0036D | 2012 | Opossums | *C. felis* | 3 positive fleas / 597 fleas | [22] |
| Taiwan | Isolate HA0985 | 2012 | Cat | *C. felis* | ND | Genbank (accession: MZ773249) |
| Brazil | Isolate 19P | 2013 | Opossums | *C. felis* | 1 positive flea / 54 fleas | [23] |
| Australia | Isolate AL357-5 | 2014 | Cat | *C. felis* | 1 positive flea / 11 fleas | [24] |
| Kazakhstan | ND | 2014 | *Rhombomys opimus* | *Coptopsylla lamellifer* | 374 fleas – 1 positive pool / 40 pools | [25] |
|  |  |  |  | *Xenopsylla skrjabini* | 1,444 fleas – 1 positive pool / 73 pools |  |
|  |  |  |  | *Xenopsylla gerbilli* | 532 fleas – 6 positive pools / 87 pools |  |
| India | Isolate RAP2CF15 | 2015 | Dog | *C. felis* | 4 positive fleas / 16 fleas | [26] |
| Marianna Islands, USA | ND | 2015 | *Rusa marianna* | *Rhipicephalus microplus* | 1 positive tick / 78 ticks | [27] |
| Oklahoma, USA | ND | 2016 | Privately-owned cat | *C. felis* | 222 fleas – 1 positive pool / 72 pools | [28] |
| Chile | ND | 2016 | Rural dogs (negative PCR) | *C. felis* | 126 fleas – 1 positive pool / 68 pools | [29] |
| Colombia | Isolate P227 | 2017 | *Equus caballus* | *Dermacentor nitens* | 2,809 ticks – 1 positive pool / 602 pools | [30] |
| Cambodia | ND | 2017 | *Canis familiaris* | *C. orientis* | 2 positive fleas / 126 fleas | [31] |
| Israel | Isolate CtlS-18 | 2017 | Cat | *C. felis* | 2 positive fleas / 91 fleas | [32] |
| Texas, USA | Isolate F112 | 2017 | *Felis catus* | *C. felis* | 721 fleas – 28 positive pools / 140 pools | [33] |
|  |  |  |  | *Felis catus* | 1 positive cat / 24 cats |  |
| Texas, USA | Isolate 21.1 | 2017 | Cat | *C. felis* | 314 fleas – 6 positive pools / 87 pools | [34] |
| Uganda | Isolate MUWRP-0110 | 2017 | Rabbit | *C. canis* | 353 fleas – 1 positive pool / 62 pools | [35] |
| California, USA | Isolate CA05Ctfe | 2018 | Cat | *C. felis* | 77 fleas – 5 positive pools / 64 pools | [36] |
|  |  |  |  | *Felis catus* | 2 positive cats / 30 cats |  |
| Thailand | Isolate G14-447B | 2018 | Domestic animal | *C. felis* | 88 fleas - 1 positive pool / 12 pools | [37] |
| Colombia | ND | 2019 | Dog | *C. felis* | 1,242 fleas - 1 positive pool / 245 pools | [38] |
| Malaysia | Isolate DB 37 | 2019 | Cat | *C. felis* | ND | [39] |
| Thailand | Isolate THAI | 2020 | Dog | *C. orientis* | 1 positive flea / 248 fleas | [40] |
| Thailand | ND | 2021 | Dog | *C. felis* | 15 positive fleas / 95 fleas | [41] |
|  | ND | 2021 | Dog | *Echidnophaga gallinacea* | 1 positive flea / 95 fleas |  |
|  | Isolat 706A | 2021 | Dog | Tick | 1 positive tick / 3 ticks |  |
| Guatemala | ND | 2022 | Dog | *C. felis* | 1 positive flea / 433 fleas | [42] |
| India | ND | 2022 | Cat | *C. felis* | 39 fleas – 1 positive pool / 8 pools | [43] |
|  |  |  | Dog | *C. canis* | 18 fleas – 1 positive pool / 6 pools |  |
| Mexico | Isolate MTY | 2023 | Dog | *C. felis* | ND | Genbank (accession: OR651412) |
| India | Isolate G5 | 2023 | Domestic animals and rodents | *C. felis* | 90 fleas – 1 positive pool | [44] |
| Portugal | ND | 2025 | Cat | *C. felis* | 1 positive flea / 1,188 fleas | [45] |

To address this, we present the formal description of *Rickettsia senegalensis* sp. nov. based on the thorough characterization of its type strain, PU01-02ᵀ, which was successfully adapted to *in vitro* culture. This study provides the crucial biological and genomic foundation required to officially validate this taxon and enable future experimental research on this emerging pathogen.

## MATERIALS AND METHODS

### Origin and primary isolation of strain PU01-02^T^

*Rickettsia senegalensis* strain PU01-02ᵀ was originally isolated in 2012 from *C. felis* cat fleas collected from cats in Dakar, Senegal [21]. As previously described, live fleas were homogenized and inoculated into XTC-2 cell monolayers using the shell-vial centrifugation technique at 28 °C. Rickettsial infection was monitored over two weeks by Gimenez staining and confirmed by specific *gltA* quantitative PCR.

### Cell culture

Strain PU01-02^T^ of *R. senegalensis* was maintained in cell culture using monolayers of *Xenopus laevis* XTC-2 cells incubated at 28 °C in a CO₂-free atmosphere. Rickettsial cultures were maintained in 25 cm² flasks with Leibovitz’s L-15 medium (Thermo Fisher Scientific, Illkirch-Graffenstaden, France), supplemented with 5% fetal bovine serum (FBS, Thermo Fisher Scientific) and 8% tryptose phosphate broth (Sigma Aldrich Chimie Saint-Quentin-Fallavier, France). Strain PU01-02^T^ was also inoculated into *Spodoptera frugiperda* SF9 cells and *Lymantria dispar* LD652 cells, incubated at 28 °C in TC-100 medium (Thermo Fisher Scientific) supplemented with 5% FBS. Additionally, the strain was cultured in transformed mouse (*Mus musculus*) fibroblast L929 cells, incubated at 37 °C with 5% CO₂ in Dulbecco’s Modified Eagle’s Medium (DMEM, Thermo Fisher Scientific) supplemented with 5% FBS. Detection of rickettsiae was performed using Gimenez staining and quantitative PCR, following previously described protocol and primers [12].

### Molecular analyses

Initially, the sequencing of key taxonomic markers (*gltA*, *ompB*, *rrs*, and *sca4*) was performed for preliminary identification and compliance with standard rickettsial taxonomic guidelines, which was subsequently complemented and validated by whole-genome sequencing. DNA was extracted from the cell culture supernatant using the EZ1 automated system and the DNeasy Tissue Kit (Qiagen, Hilden, Germany). The *gltA* (1,250 bp), *ompB* (4,858 bp), *rrs* (1,421 bp), and *sca4* (1,677 bp) genes were sequenced using previously described primer pairs [46]. Amplicons were purified using Macherey-Nagel purification plates (Macherey-Nagel EURL, Hoerdt, France) and sequenced with the BigDye Terminator Cycle Sequencing Kit (Thermo Fisher Scientific). Sequencing reactions were analyzed on an Applied Biosystems 3500 Genetic Analyzer (Thermo Fisher Scientific). The resulting sequences were assembled using ChromasPro software (version 1.7; Technelysium Pty Ltd., Tewantin, Australia). Multiple sequence alignments were performed with MUSCLE implemented in MEGA X (Molecular Evolutionary Genetics Analysis version X), and phylogenetic trees based on the *gltA* and *ompB* genes were inferred using the Maximum Likelihood method in MEGA X, applying the Kimura two-parameter model with 1,000 bootstrap replicates.

### Genomic sequencing

Following assembly, host-derived (XTC-2) contigs were bioinformatically filtered out and removed based on sequence coverage and BLASTn searches against the host database. XTC-2 cells and bacteria from five 150 cm² flasks were collected and purified following a previously established protocol [47]. After a two-hour enzymatic lysis with proteinase K at 56°C, genomic DNA was extracted using the EZ1 DNA Tissue Kit on the EZ1 automated system (Qiagen, Hilden, Germany). DNA quantification was performed using the Qubit High-Sensitivity Assay (Thermo Fisher Scientific). Genomic DNA was sequenced using MiSeq technology (Illumina Inc., San Diego, CA, USA) and the GridION platform (Oxford Nanopore Technologies, Oxford, UK), as previously described [48].

Reads obtained from Illumina MiSeq and Nanopore sequencing were assembled using Unicycler [49]. Following assembly, host-derived (XTC-2) contigs were bioinformatically filtered out and removed based on sequence coverage and BLASTn searches against the host database. OrthoANI (Orthologous Average Nucleotide Identity) was used to evaluate the genomic similarity between strain PU01-02^T^ and the species with validly published names of the TRG, applying a species delineation threshold of 99.19% [50]. Furthermore, the Genome-to-Genome Distance Calculator (GGDC) 2.1 web server (https://tygs.dsmz.de/) was utilized to determine the digital DNA-DNA hybridization (dDDH) values between the selected genomes. A threshold of less than 92.30% using formula 2 was applied to define species boundaries and a genomic tree was constructed with TYGS [51].

### Electron microscopy

For ultrastructural observation, infected XTC-2 cells underwent fixation using 2.5% glutaraldehyde in a 0.1 M sodium cacodylate buffer at 4 °C for 5 h. Resin embedding was executed with a PELCO BiowavePro+ (Ted Pella, Redding, USA) microwave device. The specimens were then sequentially dehydrated through an increasing gradient of ethanol (50%, 70%, and 96%) prior to infiltration with medium-grade LR-White resin (Polysciences, Warrington, USA). After a 72 h polymerization phase at 60 °C, 100-nm ultrathin sections were prepared with a Leica UC7 ultramicrotome and recovered onto HR25 300-mesh copper/rhodium grids (TAAB, Aldermaston, UK). Post-staining of the sections was performed using uranyl acetate and lead citrate for contrast enhancement. Finally, grids were fixed onto a glass slide with double-sided adhesive tape and coated with platinum. Specimen grids were observed by scanning electron microscopy (SEM) using a SU5000 microscope (Hitachi High-Technologies, Tokyo, Japan) operated under high vacuum at an accelerating voltage of 7 kV, in observation mode with a spot size 30 and a backscattered electron (BSE) detector, following a previously described protocol [52].

## RESULTS

### Primary isolation

Primary isolation from *C. felis* fleas collected in Dakar, Senegal yielded three rickettsial strains (PU01-02, PUX1-X2, and PU03-04) in XTC-2 cells [21]. All exhibited the typical intracellular Gimenez-positive morphology. Among these isolates, strain PU01-02ᵀ was selected for the present taxonomic characterization.

### Cell culture

*Rickettsia senegalensis* was maintained on XTC-2 cell monolayers for 29 passages. The SF9 and LD652 cell lines also supported satisfactory growth of the strain PU01-02^T^ (Figure 2). After 28 days, the Ct values obtained for these three cell types were 17.05, 17.98 and 19.76 for XTC-2, SF9 and LD652 cells, respectively. In all three cell lines (XTC-2, SF9 and LD652), rickettsiae were observed both intra- and extracellularly during continuous culture, with no cytopathic effects. In contrast, no detection of the bacterium was achieved by Gimenez staining or quantitative PCR in L929 cells after 40 days post-inoculation.

**Figure 2.**
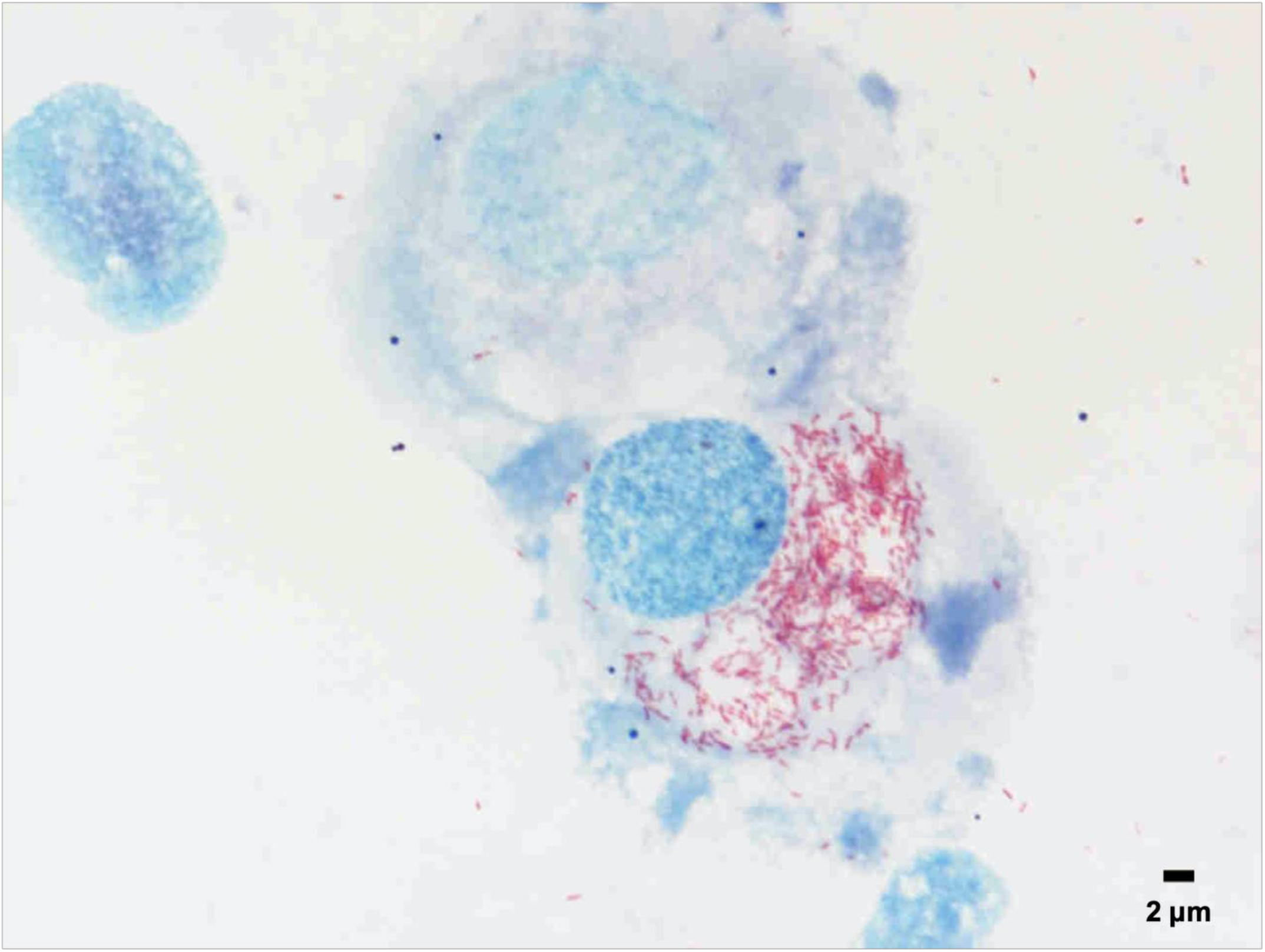
*Spodoptera frugiperda* (*Lepidoptera*) SF9 cell line infected with *Rickettsia senegalensis* sp. nov. strain PU01-02^T^, Gimenez staining, 1500x.

### Molecular identity analyses

BLAST sequence comparison for the *gltA* gene revealed that the strain PU01-02^T^ shares 100% similarity with *R. senegalensis* strain RAPCF15, which was identified in *C. felis* fleas collected in India. Additionally, a 99.91% similarity (1,149/1,150 bp) was found with *Rickettsia* sp. RF31, an isolate obtained from *C. canis* fleas collected from a dog in Thailand. The closest species with validly published names, *R. asembonensis* 8556D1, showed 98% (1,105/1,127 bp) similarity. For the *ompB* gene, a 99.81% (4,285/4,293 bp) sequence similarity was observed with the *Ca.* R. senegalensis Cf_US_0036D (identified in *C. felis* fleas collected from opossums in the USA), while the highest similarity among species with validly published names was found with *Rickettsia rhipicephali* strain HJ#5 (91.08%, 4,440/4,875 bp). Regarding the *sca4* gene, the BLAST results for the strain PU01-02^T^ demonstrated 100% (1,050/1,050 bp) and 97.17% (1,612/1,659 bp) sequence similarity with the *R. senegalensis* Cf_US_0036D and *R. felis* clone As3, respectively. As the sequence similarity values for all three genes fall below the established thresholds for defining a *Rickettsia* species, these results support the conclusion that strain PU01-02^T^ represents a new species [53].

### Phylogenetic analyses

Sequencing of the *gltA* and *ompB* genes enabled the phylogenetic analysis of *R. senegalensis* (Figure 3). The phylogenetic trees showed that the strain PU01-02^T^, along with the other isolates of *R. senegalensis*, is positioned within the TRG.

**Figure 3.**
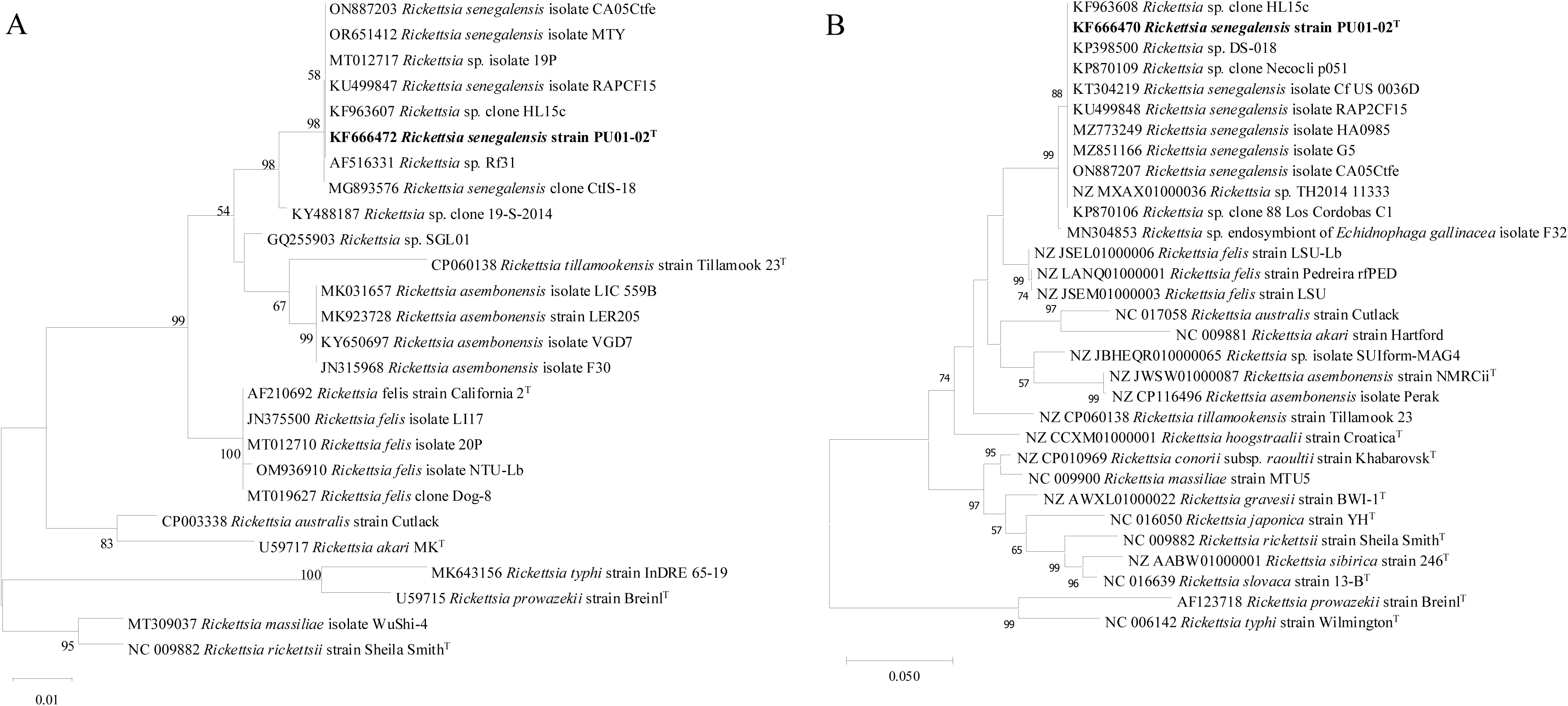
Phylogenetic trees of the *gltA* (A) and *ompB* (B) genes for *Rickettsia senegalensis* strain PU01-02^T^ and its closest species. Phylogenetic trees were reconstructed using the Maximum Likelihood method under the Kimura two-parameter model, with branch support assessed by 1,000 bootstrap replicates. Total length 1,250 bp for *gltA* and 4,858 bp for *ompB*.

### Genomic analyses

Figure 4A displays a genomic tree of the strain PU01-02^T^. This strain harbors a circular chromosome with an estimated size of 1,626,546 bp and a G+C content of 33.2%, consistent with previously reported values for other rickettsiae [54]. Genome annotation identified 1,986 protein-coding genes, along with 33 tRNA genes and three rRNA operons. The strain PU01-02^T^ shows the highest dDDH value with *R. felis* strain Pedreira, reaching 65.30%, while the maximum OrthoANI value is 95.95% with the same strain (Table S1, Figure 4B). Both values fall below the established species delineation thresholds, confirming that *R. senegalensis* represents a novel species [55].

**Figure 4.**
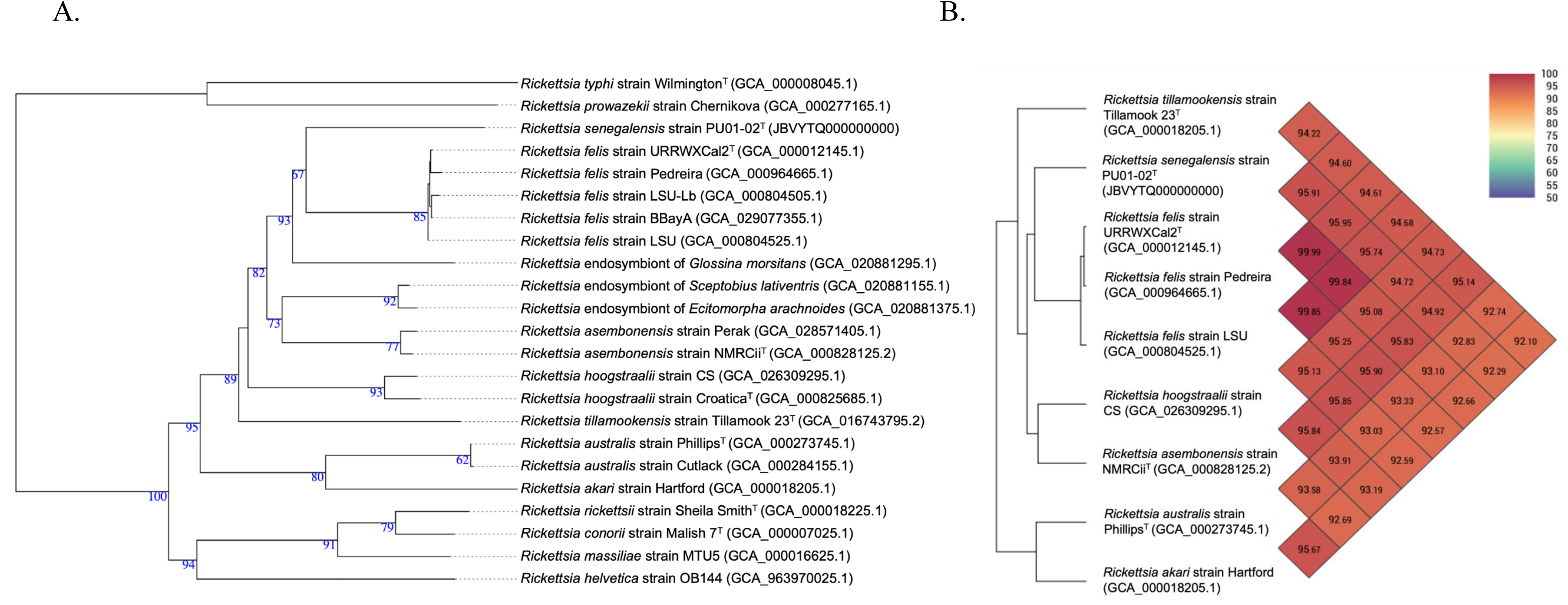
(A) Genomic tree of *Rickettsia senegalensis* strain PU01-02^T^, and closely related species constructed using TYGS. (B) OrthoANI values calculated using genome sequences of *R. senegalensis* strain PU01-02^T^ and TRG *Rickettsia* species with validly published names.

Additionally, three plasmid contigs, designated pRS1, pRS2, and pRS3, were detected through NGS sequencing, with respective sizes of 59,188 bp, 28,830 bp and 50,788 bp. The plasmid pRS01 appeared to be unique, sharing only 88.03% nucleotide identity over 63% coverage with the pRra2 plasmid from *Rickettsia raoultii* strain Khabarovsk. The highest similarity observed for plasmid pRS2 was 93.27%, with a limited coverage of 22%, when compared to the genome assembly of the *Rickettsia* endosymbiont of *Cantharis rufa* isolate 54716. Finally, plasmid pRS3 exhibited a maximum identity of 94.57% (42% coverage) with the genome assembly of the *Rickettsia* endosymbiont of *Pantillius tunicatus* isolate 54737, and 92.51% identity with 28% coverage when compared to the pRF plasmid of *R. felis*.

### Electron microscopy

The microscopic characterization of strain PU01-02^T^ was performed by scanning electron microscopy of infected cell cultures (Figure 5). Multiple bacteria exhibited a morphology typical of other *Rickettsia* species, appearing as rod-shaped cells approximately 1.5 µm in length and 0.3 µm in width. The bacteria were located exclusively within the host-cell cytoplasm, where they occurred either individually or in dense clusters, without evidence of vacuolar sequestration, consistent with active intracellular replication. The host-cell nucleus and cytoplasmic boundaries remained clearly distinguishable, while the bacteria were distributed throughout the cytoplasm surrounding the nucleus.

**Figure 5.**
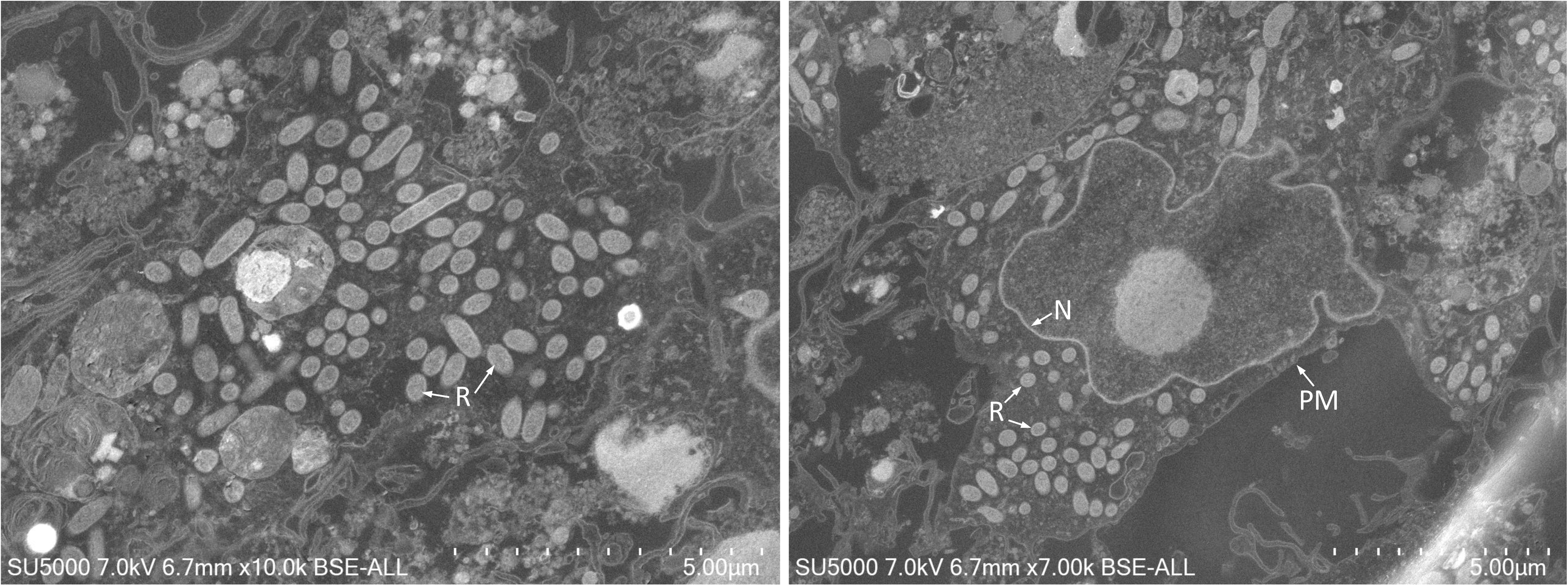
Scanning electron microscopy (SEM, BSE mode) image of *Rickettsia senegalensis* strain PU01-02^T^ infecting XTC-2 cells. R, *R. senegalensis* cells; N, host-cell nucleus; PM, plasma membrane. Scale bars = 5 µm.

## DISCUSSION

In this study, we formally describe *R. senegalensis* sp. nov., a novel *Rickettsia* species isolated in 2012 from *C. felis* cat fleas collected in Dakar, Senegal [21]. Prior to this formal taxonomic description, this bacterium was provisionally referred to as *Candidatus* Rickettsia senegalensis in the literature.

The successful isolation, stable *in vitro* cultivation, and polyphasic characterization of *R. senegalensis* strain PU01-02ᵀ represent a major milestone. Relying solely on molecular detection restricts our understanding to DNA presence, leaving the physiological and functional traits of the living bacterium unexplored. Obtaining a viable, pure isolate bypasses these limitations. Phenotypically, it allowed us to establish its unique optimal growth parameters (28°C in insect and amphibian cell lines) and ultrastructural features, demonstrating its free cytoplasmic localization without vacuolar sequestration. Genomically, having a high-quality, non-metagenomic assembly allowed for the precise detection of three distinct plasmids. Plasmids are key drivers of horizontal gene transfer and evolution within the *R. felis* cluster; thus, the viability of this strain opens the door for future experimental studies to investigate plasmid-mediated virulence, replication kinetics, and host-cell interactions under controlled laboratory conditions.

Since its first detection, this species has been identified, across all continents except Antarctica, in various hematophagous arthropods, including four flea genera (*Ctenocephalides, Coptopsylla, Xenopsylla,* and *Echidnophaga*), as well as hard tick species [22, 25, 40, 41]. Some of these arthropods were collected from domestic animals, raising concerns about potential transmission to humans [28, 37]. Although the pathogenicity of *R. senegalensis* remains unknown, its close phylogenetic relationship with *R. felis* and *R. asembonensis* (both being responsible for vector-borne diseases) suggests a possible zoonotic risk. This concern is further reinforced by the detection of *R. senegalensis* DNA in the tissues of two unowned cats captured in the Americas [36]. Previous studies have also reported *Rickettsia* DNA in the blood of dogs and cats, although it remains unclear whether the infection is sufficiently prolonged and stable to establish host competence [56, 57]. Nonetheless, these mammals play a crucial role as primary hosts for fleas, facilitating bacterial amplification through horizontal transmission.

Moreover, *R. senegalensis* has been detected in fleas collected from rodents in both the Americas and India [22, 44]. This finding underscores the need for further investigation into the role of rodents in the epidemiological cycle of TRG species. This issue is particularly relevant as rodents, the most diverse and abundant mammalian order, are already well known as both carriers and reservoirs of numerous human pathogens [58, 59]. The major host and vector appears to be *C. felis*, a flea species with a global distribution. However, most detections of *R. senegalensis* have been reported in tropical and subtropical regions.

These findings highlight the importance of enhancing surveillance efforts not only on hematophagous arthropods but also on domestic animals, due to the high zoonotic risk associated with their close contact with humans. Additionally, monitoring wild animals is crucial to gaining a better understanding of the epidemiology of TRG Rickettsiae. Although cat and dog fleas, *C. felis* and *C. canis*, appear to be the primary arthropod hosts, the ecology and distribution of these bacteria remain largely unknown [9]. Furthermore, *R. senegalensis* appears to be sympatric with other TRG species, such as *R. felis* and *R. asembonensis* [22]. Finally, formally describing *R. senegalensis* sp. nov. and depositing its type strain PU01-02ᵀ (= DSM 28250ᵀ = CSUR R184ᵀ) transitions this taxon from a provisional “*Candidatus*” status to an officially validated species. Making this fully characterized, viable reference strain publicly available is essential for the scientific community. It provides researchers with the physical material needed to perform comparative studies, avoid diagnostic misidentification within the complex *R. felis* cluster, and accurately track the global distribution and clinical relevance of this emerging *Rickettsia*.

### Description of Rickettsia senegalensis sp. nov

*Rickettsia senegalensis* (se.ne.ga.len′sis. N.L. fem. adj. *senegalensis* from Senegal, the country in western Africa where the *Ctenocephalides felis* fleas were collected) is an obligate intracellular, Gram-negative bacterium that proliferates in *Xenopus laevis* XTC-2, *Spodoptera frugiperda* SF9, and *Lymantria dispar* LD652 cell lines at 28 °C in a CO₂-free atmosphere. The bacterium does not grow at 37 °C in *Mus musculus* L929 fibroblast cultures. Phylogenetic analysis based on *gltA*, *ompB* and *sca4* gene sequences demonstrates that *R. senegalensis* is distinct from other *Rickettsia* species with validly published names. The PU01-02^T^ strain clusters with other *R. senegalensis* strains identified in various flea species. The genome size is 1.62 Mb, with a G+C content of 33.2%. The genome accession number is JBVYTQ000000000, and the plasmid accession numbers are PZ272915, PZ272916, and PZ272917, for pRS01, pRS02 and pRS03, respectively. The sequences corresponding to the *rrs*, *gltA*, *ompB*, and *sca4* genes of the PU01-02^T^ strain have been deposited in GenBank under the accession numbers KF666476, KF666472, KF666470, and KF666474, respectively.

The type strain, PU01-02ᵀ (= DSM 28250ᵀ = CSUR R184ᵀ), was isolated from a *C. felis* flea collected in Dakar, Senegal, in 2012.

## Authors and contributors

Conceptualization: CL and OM. Data curation: CL, CTH and OM. Formal analysis: CL, CTH and OM. Investigation: CL and OM. Methodology: CL and OM. Project administration: FF and OM. Resources: CL and OM. Software: CL and CTH. Supervision: FF and OM. Validation: CL, PEF, FF and OM. Visualization: CL and OM. Writing-Original draft: CL. Writing-review & editing: all authors reviewed and edited the manuscript.

## Acknowledgments

The authors thank Hitachi High-Tech Corporation (Japan) for the installation of a SU5000 microscope at IHU Mediterranean Infection.

## Supplementary material

Table S1. Genomic similarity analysis of *Rickettsia senegalensis* strain PU01-02^T^.

